# Precision Targeting of the Plasminogen Activator Inhibitor-1 Mechanism Increases Efficacy of Fibrinolytic Therapy in Empyema

**DOI:** 10.1101/2020.10.05.325688

**Authors:** Galina Florova, René A. Girard, Ali O. Azghani, Krishna Sarva, Ann Buchanan, Sophia Karandashova, Christian J. DeVera, Danna Morris, Mignote Chamiso, Kathleen Koenig, Douglas B. Cines, Steven Idell, Andrey A. Komissarov

## Abstract

Plasminogen activator inhibitor-1 (PAI-1) is an endogenous irreversible inhibitor of tissue-type (tPA) and urokinase (uPA) plasminogen activators. PAI-1-targeted fibrinolytic therapy (PAI-1-TFT) is designed to decrease the therapeutic dose of tPA and uPA to attenuate the risk of bleeding and other complications. The docking site peptide (DSP) is a part of the PAI-1 reactive center loop, which interacts with plasminogen activators, thus affecting the PAI-1 mechanism. We used DSP for PAI-1-TFT in two rabbit models: chemically-induced pleural injury and *Streptococcus pneumoniae* induced empyema. PAI-1-TFT with DSP combined with single chain uPA or tPA resulted in an up to 8-fold decrease in the minimal effective therapeutic dose of plasminogen activator and induced no bleeding. An increase in the level of PAI-1 in infectious pleural injury, when compared to chemically-induced injury, coincided with an increase in the minimal effective dose of plasminogen activator and DSP. PAI-1 is a valid molecular target in *S. pneumoniae* empyema model in rabbits, which closely recapitulates key characteristics of empyema in humans. Low dose PAI-1-TFT is a novel precise interventional strategy that may improve fibrinolytic therapy of empyema in clinical practice.

## Introduction

Plasminogen Activator Inhibitor-1 (PAI-1) is a mechanism-based inhibitor of tissue type (tPA) and urokinase (uPA) plasminogen activators (PAs) (1-3) used for fibrinolytic therapy. Levels of PAI-1 increase by up to three orders of magnitude in empyema (pleural sepsis), and organizing pleural injury (4-6), limiting endogenous and exogenous plasminogen activating activity that could help prevent or treat uncontrolled fibrin deposition induced by inflammation. Previously, we suggested that PAI-1 is a molecular target in treatment of chemically (tetracycline; TCN)-induced pleural injury in rabbits (7). PAI-1-targeted fibrinolytic therapy (PAI-1-TFT) is designed to affect the PAI-1 mechanism in order to decrease the minimal effective dose (MED) of a plasminogen activator. We used a novel model of pleural injury induced by *Streptococcus pneumoniae* (8) to recapitulate the inflammation and high levels of PAI-1 associated with the development of empyema in humans and its progression from the early, acute stage to more advanced chronic stages (8). Both single chain (sc) tPA and scuPA in a bolus MED 2.0 mg/kg are effective in clearing the pleural space in rabbits with acute *S. pneumoniae* empyema (8). The MED for empyema is 4- and 13.8-fold higher than MEDs of scuPA and sctPA in TCN-induced pleural injury in rabbits (9). This increase in the MED in *S. pneumoniae* empyema could reflect changes in the molecular mechanism of intrapleural fibrinolysis due to: (i) a slower rate of fibrinolysis, (ii) faster *in vivo* inactivation of plasminogen activators, (iii) an increase in the intrapleural level of PAI-1, or a combination of these factors. We reasoned that if PAI-1 is a true molecular target for pleural injury then PAI-1-TFT should reduce the MED of plasminogen activators in both chemical and infectious pleural injury, and chose to target the docking site (DS) of PAI-1. uPA and tPA with alanine mutations of positively charged residues in the 37-loop, which participates in exosite interactions (10;11) (ΔDS-uPA and tenecteplase, respectively), react with PAI-1 at a rate that is almost two orders of magnitude slower than the WT enzymes (12-14). Fibrinolytic therapy with ΔDS-scuPA in TCN-induced pleural injury in rabbits showed a trend towards improved outcomes compared to WT-scuPA (14), indicating that targeting DS interactions could increase the efficacy of fibrinolytic therapy, although its success was limited by its short intrapleural half-life. These results provide a rationale to further explore precision targeting of DS interactions with a PAI-1 derived Docking Site Peptide (DSP; EEIIMD), which was used successfully in other translational research (15;16). Targeting PAI-1 mechanism with mAbs that redirect the reaction with plasminogen activator towards the substrate branch (17) or accelerate active to latent transition of PAI-1 (14) resulted an in up to 8-fold decrease in the MED of scuPA in treatment of TCN-induced pleural injury. Previously, we hypothesized that DSP-mediated PAI-1-TFT also will increase the efficacy of fibrinolytic therapy (14). To test this hypothesis, we evaluated the effect of DSP, a small peptide molecule, that unlike anti-PAI-1 mAbs interacts with the enzyme, on the outcomes of fibrinolytic therapy with sctPA and scuPA in rabbit models of chemically-induced and infectious pleural injury.

## Results

### Developing and treating chemically-induced and infectious pleural injury in rabbits

Two validated models of pleural injury (chemically-induced and infectious, Figure 1) (7-9;14;17-19) were used to evaluate the effect of DSP on the outcomes of fibrinolytic therapy. Pleural injury was monitored and verified using ultrasonography. Levels of PAI-1 (active and total), TGF-β, TNF-α, IL-6 and IL-8 (Figure 1A-F) were measured in samples of pleural fluid collected immediately prior to PAI-1-TFT (at 48 or 72h for the chemically-induced and infectious injuries, respectively). Notably, while levels of PAI-1, TGF-β, TNF-α, IL-6, and IL-8 were elevated in both models, they were higher (*P*<0.05) in empyema (Figure 1A-F). Both models featured robust intrapleural fibrin deposition detected by ultrasonography (data not shown) and documented by photography during necropsy (Figure 1G,I). The highest Gross Lung Injury Score (GLIS=50), which corresponds to fibrin strands, nets, and aggregates that were too numerous to count (TNTC), was the same for both models (Figure 1G,I). While infectious pleural injury caused greater fibrin deposition on visual inspection (Figure 1I), there was no statistically significant difference in the pleural thickening (Figure 1H,J). Thus, using DSP for PAI-1-TFT in models of pleural injury that feature different levels of PAI-1 and inflammatory responses (Figure 1), allows us to validate the approach, assess the effects of increases in the molecular target on doses of plasminogen activator and DSP, and test the concept that precision PAI-1-TFT improves therapeutic outcomes.

**Figure 1.**
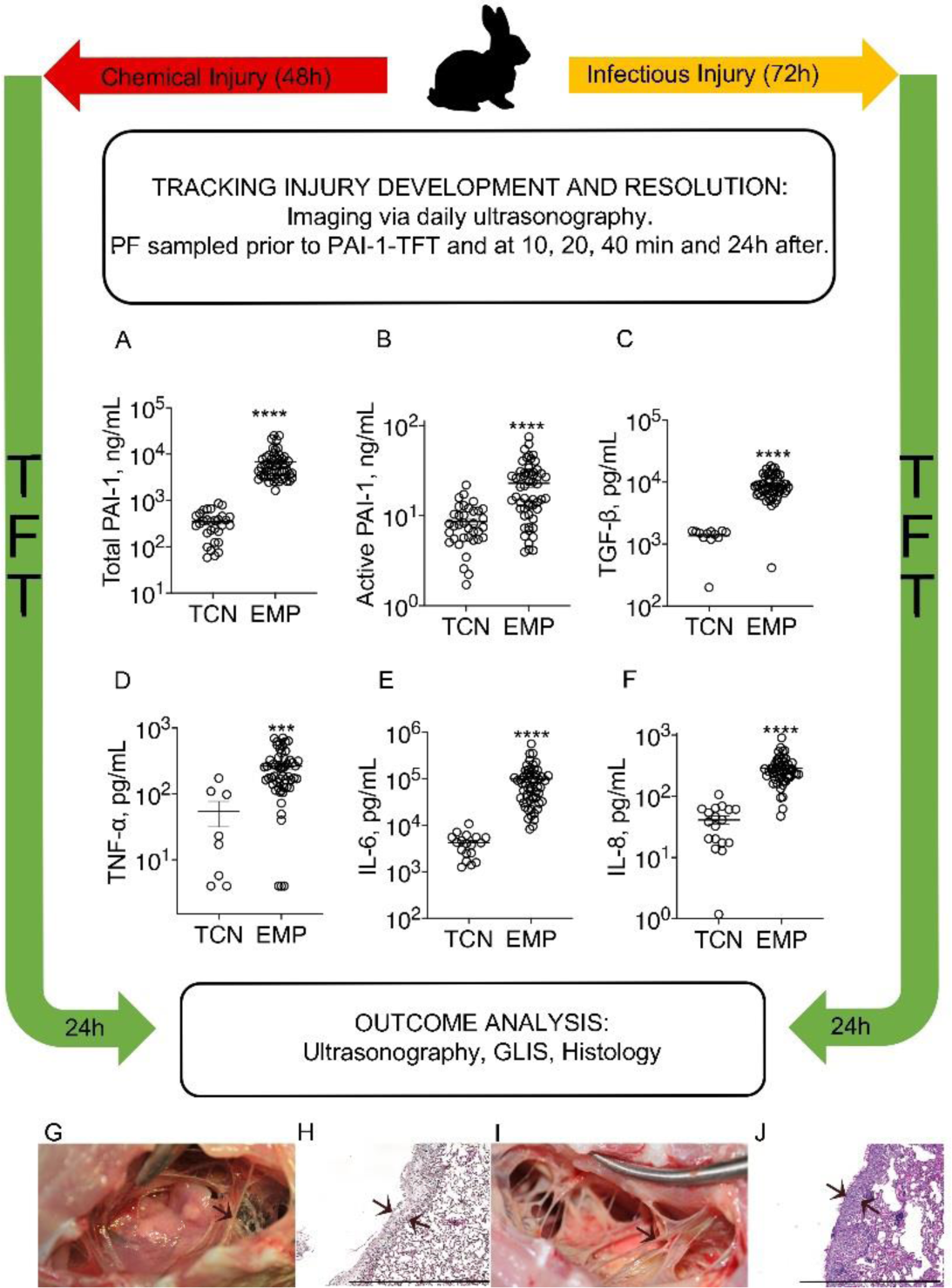
Models of chemically-induced and infectious pleural injury in rabbits were used to test PAI-1-TFT. Time course of initiation, development, of TCN-(left; 0-48h) and *S. pneumoniae* (right; 0-72h) induced pleural injury in rabbits followed by treatment (TFT), and data analysis. Pleural injury was initiated by intrapleural injection of either TCN (9) or *S. pneumoniae* (8). Injury progression was tracked using ultrasonography. Intrapleural levels of PAI-1, total (**A**; n=30 and 55) and active (**B**; n=36 and 55); TGF-ß (**C**; n=12 and 55); TNF-α (**D**; n=8 and 55); IL-6 (**E**; n=18 and 55); IL-8 (**F**; n=18 and 53), were determined in baseline samples of pleural fluid (PF) collected from animals with chemically-induced (TCN) and infectious (EMP) pleural injury, respectively. Data are presented as scatter plots, showing mean with SEM, *P*<0.01 (**), *P*<0.0001 (****) using a two-tailed Kolmogorov-Smirnov test. PAI-1-TFT was delivered at 48 (chemically-induced) and 72 (infectious) hours and outcomes were assessed 24h later using ultrasonography (not shown), postmortem visualization of the pleural space during necropsy, and by hematoxylin and eosin (H&E) of tissue samples. A Gross Lung Injury Score (GLIS) (8;17) was used for visual assessment; GLIS ranges from 0 to 50, where 0 refers to a clear pleural space and 50 to fibrin formations that are too numerous to count (TNTC); (GLIS≤10 considered successful PAI-1-TFT. Representative photographic images of GLIS=50 (arrows indicate fibrin adhesions) for TCN-induced (**G)** and infectious (**I**) pleural injury after treatment with a vehicle control. Representative images of H&E staining of paraffin-embedded lung tissue demonstrate pleural thickening in TCN-induced (**H**) and infectious (**J**) injury. Arrows indicate the pleural surface and the basement membrane of the pleura. There was no statistically significant difference (*P*>0.05) in pleural thickening between models. Images were obtained at 4x on a Cytation 5 (Biotek) with 5-6 rabbits/group; scale bars, 1 mm.

### DSP improves therapeutic outcomes of ineffective doses of plasminogen activators in both chemically-induced and infectious pleural injury in rabbits

The treatment and outcomes analyses of the two models of pleural injury are shown in Figure 1. First, DSP alone was used to treat pleural injury in both models to determine its effects on the endogenous fibrinolytic system. The results (Figure 2) clearly indicate that even a relatively high dose of DSP (2.0 mg/kg) is insufficient to mobilize endogenous fibrinolysis effectively once pleural injury has developed; there was no reversal effect with DSP alone. DSP alone, even at a 4-fold higher dose (8.0 mg/kg), was also ineffective (GLIS>10) in treatment of empyema (Figure 2). Next, PAI-1-TFT with a bolus injection of sctPA or scuPA in combination with DSP was delivered at 48h (Figure 2A; TCN) and at 72h (; Figure 2A; Empyema) after initiation of pleural injury (Figure 1). Control animals were treated with plasminogen activator alone (Figure 2). Aliquots of pleural fluid were withdrawn prior to treatment and at 10, 20, 40 min and at 24h after PAI-1-TFT. Outcomes were assessed 24h after treatment using ultrasonography and morphometric GLIS scores determined at necropsy. PAI-1-TFT was considered successful if GLIS≤10, as we previously reported (7;17). To test the effect of DSP on the outcome in TCN-induced pleural injury, ineffective (GLIS>10) doses of scuPA and sctPA were selected. The dose of scuPA (62.5 µg/kg; 1/8 of the MED for scuPA alone (9) previously shown to be effective when used with anti-PAI-1 mAbs for treatment of TCN-induced injury (14;17), and an equimolar dose of sctPA (72.5 µg/kg; 1/2 of the MED for sctPA alone (9)). The results of DSP-mediated PAI-1-TFT in chemically-induced pleural injury are shown in Figure 2A, TCN. DSP (2.0 mg/kg) renders otherwise ineffective doses of both scuPA and sctPA effective (GLIS≤10; Figure 2B,C). Notably, PAI-1-TFT caused a statistically significant (*P*<0.01) decrease in pleural thickening (Figure 2F,G versus Figure 1H). Targeting exosite interactions between PAI-1 and plasminogen activator using DSP resulted in a 2- and 8-fold increase in the efficacy of fibrinolytic therapy with sctPA and scuPA, respectively (Figure 2A, TCN). *S. pneumoniae* induced pleural injury results in higher levels of the molecular target, PAI-1, compared to TCN-induced injury (Figure 1A,B). Moreover, MEDs of sctPA (0.145 mg/kg) and scuPA (0.5 mg/kg) that effectively treat of chemically-induced injury were ineffective in empyema (8). The increase in intrapleural PAI-1 in infectious injury (Figure 1) correlates with an increase in the MED to 2.0 mg/kg (9) for both sctPA and scuPA. A dose of 0.5 mg/kg, ineffective for both sctPA (n=6) and scuPA (n=6) (Figure 2A, Empyema), was selected to test the outcomes of DSP-mediated PAI-1-TFT. First, 2.0 mg/kg DSP, a dose that was effective in chemically-induced pleural injury (Figure 2A, TCN) in combination with 0.5 mg/kg of sctPA or scuPA, was tested. In contrast to the treatment of chemically-induced injury (Figure 2A, TCN), 2.0 mg/kg DSP adversely affected the outcome of fibrinolytic therapy with scuPA but showed a trend (*P*>0.05) towards improvement with sctPA (Figure 2A, Empyema). An increase in the level of the molecular target, PAI-1, that results in a higher MED for plasminogen activator in the model of infectious pleural injury suggests that more DSP is needed to improve the outcome of PAI-1-TFT. This hypothesis was tested by sequential two-fold escalations of the DSP dose in combination with sctPA (0.5 mg/kg) (Figure 2A, Empyema). An increase in the dose of DSP from 2.0 to 4.0 (n=5) and 8.0 mg/kg (n=6) (Figure 2A, Empyema) led to effective (GLIS≤10) PAI-1-TFT with sctPA (Figure 2E). There was no statistically significant decrease in pleural thickening after successful PAI-1-TFT (Figure 2I) compared to vehicle controls (Figure 1J). These results show that an increase in the level of the molecular target in empyema correlates with an increase in the MED of sctPA and a corresponding increase in the dose of DSP needed for effective PAI-1-TFT. In contrast to sctPA, combining DSP (2.0 and 8.0 mg/kg) with scuPA (0.5 mg/kg) was less effective than scuPA alone (Figure 2A, Empyema), resulting in GLIS>10 and robust pleural thickening (Figure 2D,H). Thus, DSP, even at a high dose such as 8.0 mg/kg, affects tPA and uPA induced intrapleural fibrinolysis differently in our model of infectious pleural injury in rabbits. Next, we determined whether the MED of sctPA in empyema treatment could be further reduced when administered with 8.0 mg/kg DSP. Animals with acute empyema were treated with sctPA (0.25 mg/kg) alone (n=6) and in combination with 8.0 (n=6) mg/kg DSP (Figure 3). Clearance of fibrin from the pleural space was visualized by ultrasonography (Figure 3B,E) and confirmed using photography (Figure 3C,F). DSP (8.0 mg/kg) dramatically improves the outcome of PAI-1-TFT (*P*<0.05), converting an ineffective dose of sctPA 0.25 mg/kg to a MED (Figure 3A,C,F) that is 8-fold less than the MED of sctPA alone (2.0 mg/kg; (8)). There was no statistically significant difference in pleural thickness between successful (Figure 3G) and unsuccessful (Figures 1J, 3D) treatments.

**Figure 2.**
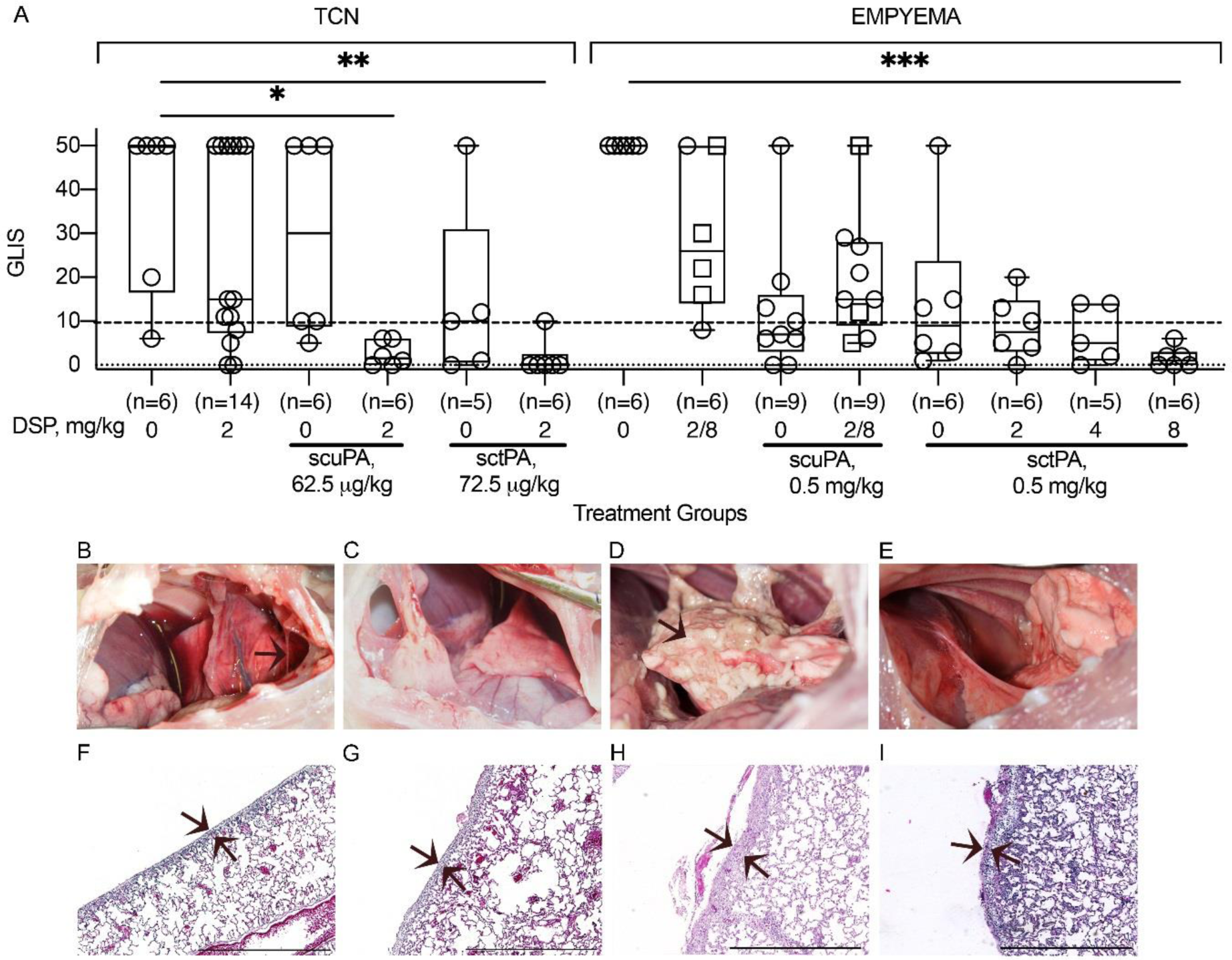
PAI-1-TFT results in a decrease in the dose of plasminogen activator in chemically-induced (TCN) and infectious (empyema) pleural injury. The efficacy of fibrinolytic therapy was measured by GLIS (GLIS≤10, at or below the dashed line, indicates successful PAI-1-TFT). **(A; TCN)** Animals were treated (from left to right) with: vehicle control (n=6); 2.0 mg/kg DSP alone (n=14); an ineffective dose (62.5 µg/kg) of scuPA (n=6); 62.5 µg/kg of scuPA with 2.0 mg/kg DSP (n=6); an ineffective dose (72.5 µg/kg) of sctPA (n=5); 72.5 µg/kg of sctPA with 2.0 mg/kg DSP (n=6); **(A; empyema)** Animals were treated (from left to right) with: vehicle control (n=6); 2.0 (n=2) or 8.0 (n=4; squares) mg/kg DSP alone; ineffective dose of scuPA (0.5 mg/kg; n=9); 0.5 mg/kg of scuPA with 2.0 mg/kg (n=6) or 8.0 mg/kg (n=3; squares) DSP; ineffective dose (0.5 mg/kg) of sctPA alone (n=6) and with 2.0 (n=6), 4.0 (n=5), and 8.0 mg/kg (n=6) DSP. Statistically significant differences determined using Kruskal-Wallis test; *, ** and *** denote *P*<0.05; <0.01 and <0.001, respectively. Representative photographic images of pleural spaces of animals with TCN-induced pleural injury successfully (GLIS=0) treated with 2.0 mg/kg DSP with 62.5 µg/kg of scuPA (**B**); or 72.5 µg/kg of sctPA (**C**) and animals with infectious pleural injury treated with 8.0 mg/kg DSP with 0.5 mg/kg scuPA (GLIS=50; **D**) or with 0.5 mg/kg sctPA (GLIS=0; **E**). Representative images of H&E staining of paraffin embedded lung tissue from the same animals (**F-I**). While animals with TCN-induced injury (**F, G**) demonstrate a statistically significant decrease in pleural thickening (*P*<0.01) compared to vehicle treated controls (Figure 1), animals with infectious pleural injury (**H, I**) did not (*P*>0.05). Statistical significance was determined using an unpaired, 2-tailed Kolmogorov-Smirnov test. Scale bars, 1mm. Images were obtained at 4x on a Cytation5 (Biotek) with 5-6 rabbits/group.

**Figure 3.**
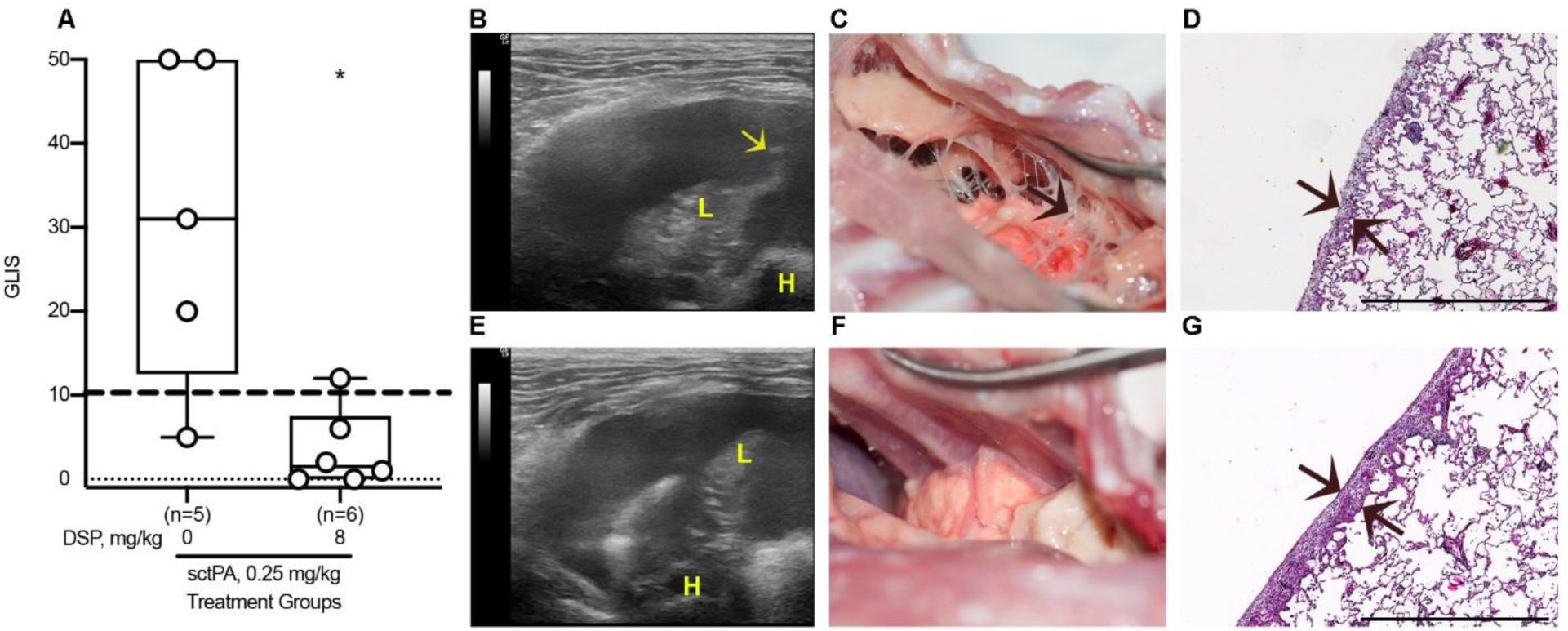
DSP (8.0 mg/kg) increases the efficacy of intrapleural fibrinolytic therapy by 8-fold and converts an ineffective dose of sctPA (0.25 mg/kg) into a MED for treatment of acute empyema in rabbits. The effect of DSP on the efficacy of fibrinolytic therapy of infectious pleural injury with an ineffective dose of sctPA (0.25 mg/kg), measured by GLIS (GLIS≤10 at or below the dashed line indicates successful PAI-1-TFT (**A**). Animals with acute empyema were treated (from left to right) with sctPA (0.25 mg/kg) alone (n=5) or together with 8.0 mg/kg DSP (n=6). The data are shown as a box plot; * denotes a statistically significant (*P*<0.05) difference determined using an unpaired, 2-tailed Kolmogorov-Smirnov test. Representative ultrasonography images of the pleural space 24h after treatment with 0.25 sctPA alone (**B**), and in combination with 8.0 mg/kg DSP (**E**) show an acoustic window for lung (L), heart (H), and a fibrin strand (arrow) in the pleural space. Photographs of the pleural space 24h after unsuccessful (GLIS=50; arrow indicates fibrin adhesions) treatment with sctPA alone (**C**), and successful fibrinolytic therapy (GLIS=5) with sctPA and DSP (**F**). H&E staining of paraffin embedded lung tissue of animals treated with sctPA alone (**D**) and together with DSP (**G**). Arrows indicate the pleural surfaces and basement membranes of the pleura. Scale bars: 1 mm. Images were obtained at 4x on a Cytation 5 (Biotek) with 5 rabbits/group.

### Pharmacokinetics of DSP in the rabbit pleural space

The results of Liquid Chromatography in tandem with Mass Spectrometry (LC MS/MS) analyses demonstrated an increase in the level of DSP in pleural fluid as the dose of the peptide was increased from 2.0 to 8.0 mg/kg (Figure 4). The intrapleural level of DSP decreases sharply in the first 10 min after injection (not shown), followed by a slower phase with apparent first order rate constants of DSP clearance (k_clr_) ranging at 0.03-0.07 min^-1^ in both models. Pharmacokinetic analyses of DSP in pleural fluid of animals with TCN-induced pleural injury (Figure 4A) and empyema (Figure 4B) confirm the presence of DSP for over 40 min after treatment as well as an increase in the intrapleural level of DSP with dose escalation (Figure 4B). The highest k_clr_ were observed in the model of TCN-induced injury (0.05±0.01, 0.06±0.01 and 0.07±0.02 min^-1^ for 2.0 mg/kg DSP alone and together with sctPA or scuPA, respectively). The rate of DSP elimination becomes slower (*P*<0.05) in rabbit model of empyema (k_clr_ were 0.04±0.02 and 0.05±0.02 min^-1^, for 2.0 mg/kg DSP with sctPA or scuPA, respectively). Increasing the dose of DSP to 4.0 and 8.0 mg/kg did not affect dramatically rate of the slow phase of DSP clearance. The values of k_clr_ for treatments with 0.5 mg/kg of sctPA with 4.0 and 8.0 mg/kg DSP and 0.25 mg/kg of sctPA with 8.0 mg/kg DSP were 0.03±0.02, 0.05±0.03 and 0.04±0.02 min^-1^, respectively. The k_clr_ for treatment with 8.0 mg/kg DSP alone (0.04±0.01 min^-1^) was similar to those observed with plasminogen activators. Thus, fibrinolytic therapy does not affect the rate of elimination of DSP from the pleural space in either model of pleural injury. The average intrapleural concentration of DSP 40 min after treatment with 2.0 mg/kg increases from 10 µg/mL in TCN-induced injury to 36 µg/mL in empyema, and increases further to >300 µg/mL after a dose escalation to 8.0 mg/kg DSP, with which PAI-1-TFT is successful (Figure 4; green symbols). These results indicate clearly that an increase in the efficacy of PAI-1-TFT in infectious pleural injury with otherwise ineffective doses of sctPA correlates with an increase in the intrapleural level of the peptide. Furthermore, these results support the hypothesis that an increase in the level of the molecular target, PAI-1, observed in empyema model (Figure 1) necessitates an increase in both the MED of sctPA and DSP to make PAI-1-TFT successful.

**Figure 4.**
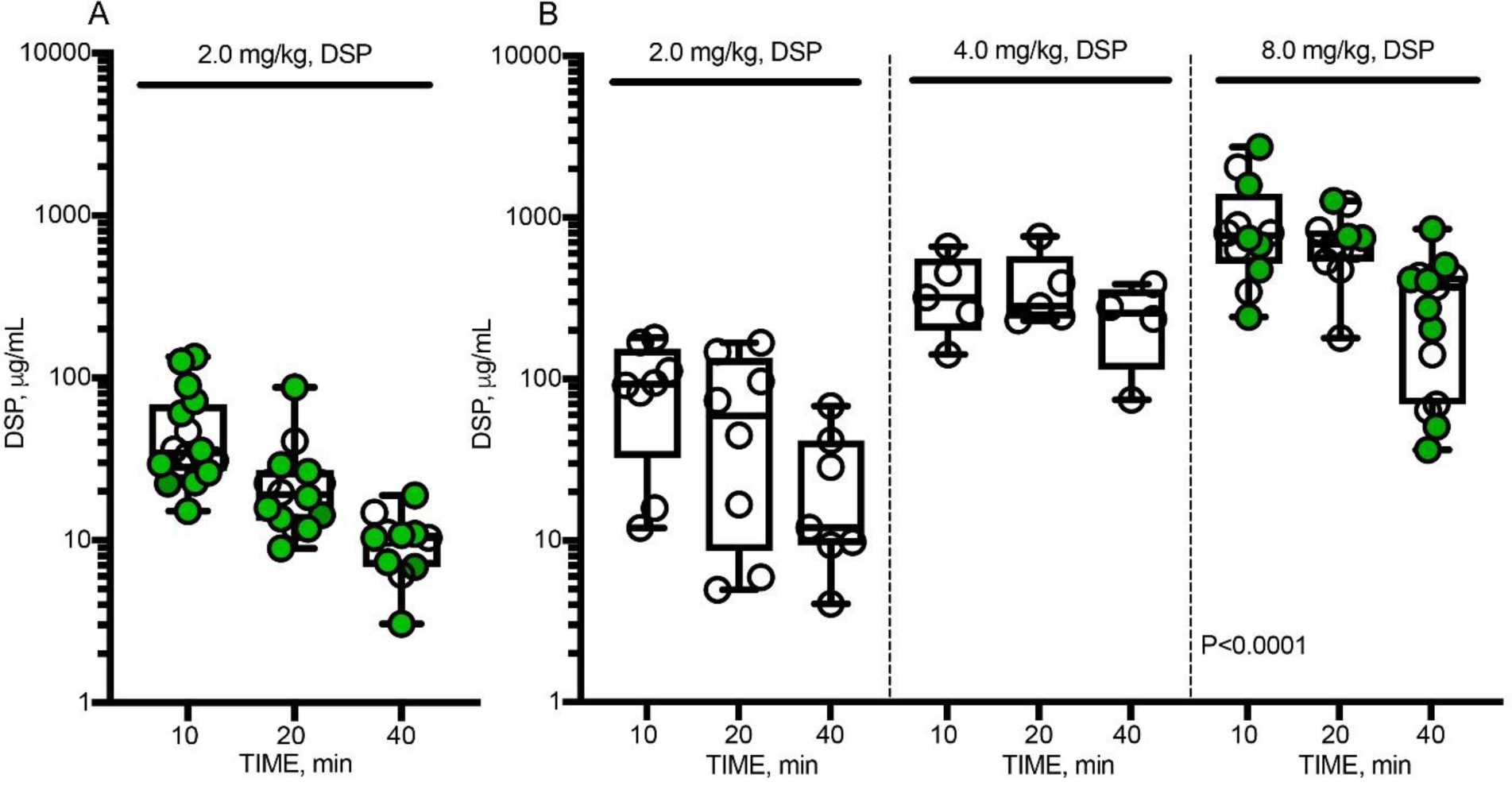
Pharmacokinetics of DSP in pleural fluid of animals with chemically-induced (A) and infectious (B) pleural injury. LC MS/MS analyses were used to measure the levels of DSP in pleural fluids of rabbits at 10, 20 and 40 min after PAI-1-TFT. Animals with chemically-induced pleural injury (**A**) were treated successfully (GLIS≤10; green circles) with DSP (2.0 mg/kg) in combination with either 72.5 µg/kg (n=6) of sctPA or 62.5 µg/kg (n=5) of scuPA; or unsuccessfully (GLIS>10; empty circles) with DSP alone (n=6); LC MS/MS analyses of pleural fluids from animals with *S. pneumoniae* induced pleural injury (**B**) treated unsuccessfully (GLIS>10; empty circles) with sctPA (0.5 mg/kg) in combination with (from left to right): 2.0 (n=6) and 4.0 (n=6) mg/kg DSP. Levels of DSP in pleural fluids of animals treated successfully (GLIS≤10; green circles) with 8.0 mg/kg DSP in combination with 0.5 (n=6) and 0.25 (n=6) mg/kg sctPA are shown in the rightmost panel. The results from unsuccessful (GLIS>10; empty circles) treatments of infectious pleural injury with DSP alone with doses of 2.0 (n=2) and 8.0 (n=4) mg/kg are added to the corresponding panels. The data are presented as box and whiskers plots, showing all points. The values of the observed first order rate constants for intrapleural DSP clearance (k_clr_), determined by fitting a single exponential equation to the data were 0.05±0.01, 0.06±0.01 and 0.07±0.02 min^-1^ (*P*<0.05, *r*>0.85) for treatment of TCN-induced injury with 2.0 mg/kg DSP alone or with 62.5 µg/kg of scuPA or with 72.5 µg/kg of sctPA, respectively (**A**), and 0.04±0.02 0.03±0.02, 0.05±0.03; 0.04±0.01, and 0.04±0.02 min^-1^, (*P*<0.05, *r*>0.6) for treatments with 0.5 mg/kg of sctPA in combination with 2.0, 4.0 and 8.0 mg/kg DSP, 8.0 mg/kg DSP alone and in combination with 0.25 mg/kg oof sctPA, respectively, (**B**).

### Changes in intrapleural plasminogen activating and fibrinolytic activities during PAI-1-TFT

Intrapleural plasminogen activating (Figure 5A-D) and fibrinolytic (Figure 5E-H) activities were measured in pleural fluids prior to (baseline) and at 10, 20 and 40 min and at 24h after treatment, as described previously (8;14;17;18). There was no detectable plasminogen activating activity at baseline or at 24h (Figure 5A-D). The fast phase of intrapleural inhibition of plasminogen activators, which occurs within the first 10 min after treatment (18), was followed by a slower phase (Figure 5A-D). The observed rate constants (k_obs_) of intrapleural inactivation during PAI-1-TFT of chemically-induced pleural injury, were in the range of 0.02-0.05 min^-1^ for tPA and uPA plasminogen activating activity. The values of k_obs_ for intrapleural inactivation of tPA (with 2.0-8.0 mg/kg DSP) and uPA (with 2.0 mg/kg DSP) during PAI-1-TFT of infectious pleural injury were 0.024±0.010 and 0.014±0.008 min^-1^, respectively. Fibrinolytic activity was suppressed at baseline and at 24h after treatment (Figure 5E-H). Supplementing pleural fluids collected at baseline and 24h with plasminogen activator (which simulates fibrinolytic therapy) resulted in a significant increase in the fibrinolytic activity (Figure 5E-H). However, over the first 10 min after PAI-1-TFT, fibrinolytic activity in pleural fluids sharply decreases until it reaches almost steady state level (Figure 5E-H), which is less than 10% of that in supplemented baseline. A rapid decrease in the pleural fluid fibrinolytic activity to a steady state level occurred with similar k_obs_ (0.40±0.11 and 0.48±0.07 min^-1^) but resulted in different steady-state levels (2.7 and 1.5%) for PAI-1-TFT with sctPA and 2.0 mg/kg DSP in chemically-induced and infectious pleural injury, respectively. Further increases in DSP (to 8.0 mg/kg in empyema treatment) did not affect k_obs_ significantly. scuPA based PAI-1-TFT with 2.0 mg/kg DSP featured a slower decrease in pleural fluid fibrinolytic activity then that observed with sctPA (0.09±0.06 and 0.16±0.06 min^-1^) and higher steady state fibrinolytic activity (5.6 and 2.5%) for chemically-induced and infectious pleural injury, respectively. The observed rate constants of intrapleural accumulation of α-macroglobulin (αM)/uPA complexes during PAI-1-TFT of chemically-induced and infectious pleural injury were 0.02±0.01 and 0.03±0.01 min^-1^, respectively (Figure 5I,K), with the maximal intrapleural concentrations of 220±80 and 260±60 nM, respectively. Both plasminogen activating (Figure 5A-D) and fibrinolytic (Figure 5E-H) activities in pleural fluid were suppressed at 24h after treatment for both successful (GLIS≤10) and unsuccessful (GLIS>10) PAI-1-TFT. No DSP was detected by LC MS/MS at 24h after any intervention (data not shown). Thus, the time needed for effective fibrinolysis (14;17) in both models of pleural injury is less than 24h, and likely similar to the one determined previously for TCN-induced pleural injury in rabbits (6-8h) (19). Notably, there was a correlation (*P*<0.05) between values of fibrinolytic potential (17) measured in pleural fluids at the baseline and at 24h after PAI-1-TFT (Figure 5J,L) indicating that fibrinolytic potential reflects status of fibrinolytic system of a specific subject. Intrapleural levels of PAI-1 (total and active), TGF-β, TNF-α, IL-6, IL-8 at 24h after PAI-1-TFT in chemically-induced and infectious pleural injury are shown in Figure 6A-F. While WBC counts in pleural fluids collected at the time of euthanasia were elevated in both rabbit models, infectious pleural injury was characterized by more than 3-fold higher total WBCs (data not shown) with a predominance of PMN cells in empyema fluids (Figure 6G). Likewise, this increase was accompanied with an increase in the markers of pleural inflammation comparable to those observed in pleural fluid prior to PAI-1-TFT (Figure 1). As shown in Figure 6, statistically significant higher total and active PAI-1 (Figure 6A,B), TGF-β, TNF-α, IL-6 and IL-8 (Figure 6C-F) were observed in pleural fluids in *S. pneumoniae*-induced pleural injury. Notably, only treatment with sctPA in combination with 8.0 mg/kg DSP led to a decrease in PMN cells (Figure 6H). However, we did not observe statistically significant changes in the levels of inflammatory markers or PAI-1 associated with successful PAI-1-TFT (Figure 6A-F, green dots). Finally, no apparent local or systemic bleeding was accompanying PAI-1-TFT in either model. There was no statistical difference in pleural fluid RBC counts in either model at any dose of DSP alone or in combination with plasminogen activator (Figure 7).

**Figure 5.**
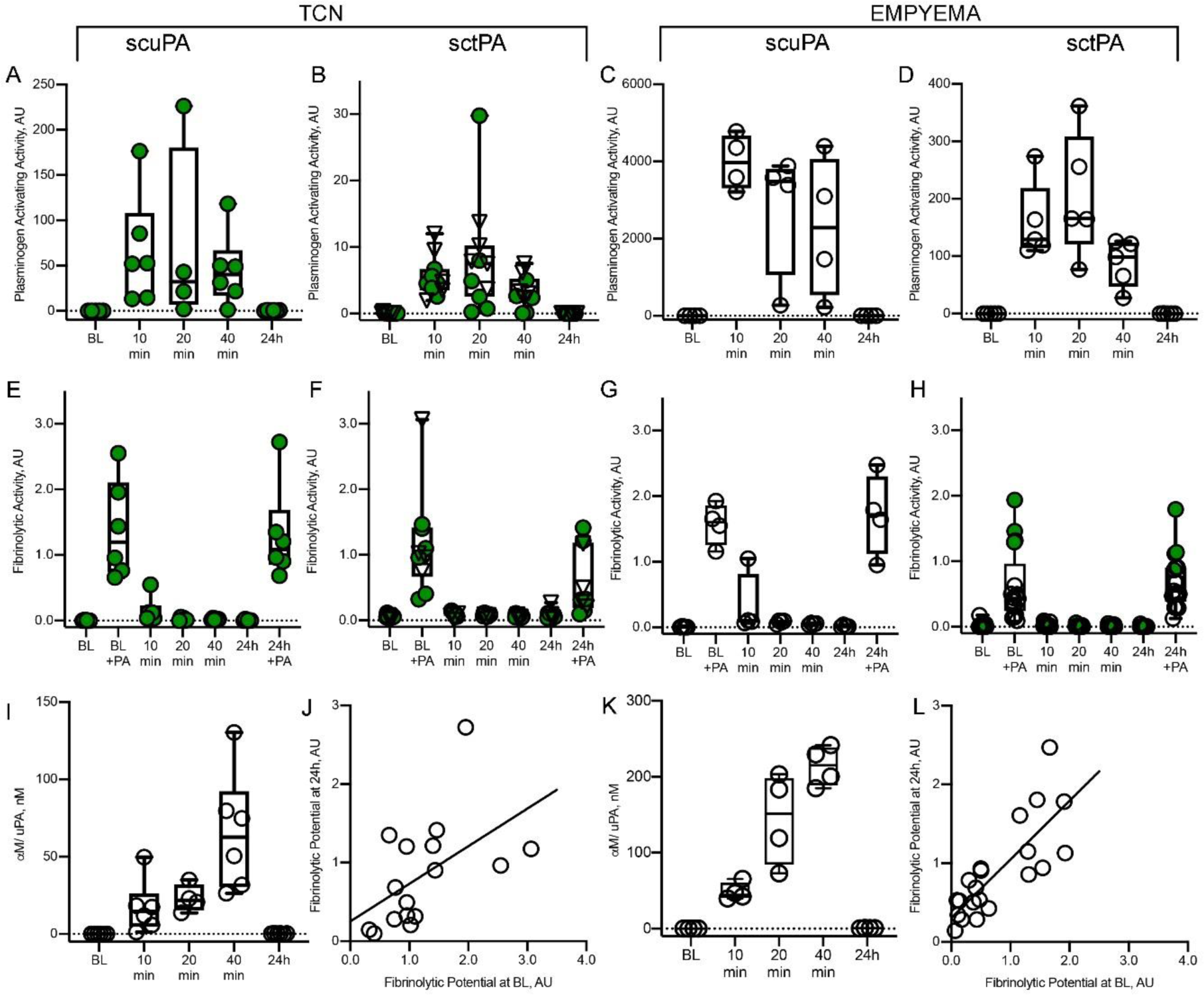
Changes in the plasminogen activating (A-D) and fibrinolytic (E-H) activities, levels of αM/uPA complexes (I, K), and correlation between fibrinolytic potential measured at the baseline and at 24h after PAI-1-TFT in pleural fluids of rabbits following chemically-induced (A,B,E,F,I,J) and infectious (C,D,G,H,K,L) pleural injury. Plasminogen activating and fibrinolytic activity was measured in samples of pleural fluid collected at baseline (BL); 10, 20, 40 min and at 24h after PAI-1-TFT with DSP 2.0 mg/kg in combination with 62.5 µg/kg of scuPA (n=6; **A, E**) or 72.5 µg/kg of sctPA (n=6; **B, F**) (chemically-induced injury; TCN); 0.5 mg/kg scuPA (n=4; **C, G**) (infectious injury; empyema). Green and empty circles represent successful (GLIS≤10) and unsuccessful (GLIS>10) PAI-1-TFT, respectively. Reversed triangles (**B, F**) represent treatment with sctPA alone (n=5, GLIS>10). **D** represents treatment with 0.5 mg/kg sctPA with 2.0 mg/kg DSP (n=6); **H**, treatments (n=17) with 0.5 mg/kg sctPA in combination with 2.0 (n=6), 4.0 (n=5) and 8.0 (n=4) mg/kg DSP and 0.25 mg/kg sctPA with 4.0 mg/kg DSP (n=2). Fibrinolytic activity in aliquots of pleural fluids withdrawn prior to (BL) and 24h after PAI-1-TFT were analyzed with (+PA) or without supplementation with 4.0 nM of plasminogen activator (17). Levels of αM/uPA in pleural fluids at BL, 10, 20, 40 min and 24h (**I, K**) were measured as described previously (18) and plotted against time. Fibrinolytic potential (17) was determined in samples of pleural fluid withdrawn at 24h and plotted versus corresponding values determined for baseline (BL) for TCN-induced injury (n=16; **J**) and empyema (n=21, **L**). The solid lines represent the best fit (*P*<0.05) of a linear equation to the data with slopes 0.48±0.20 and 0.74±0.13, and *r*=0.54 and 0.80 for TCN-induced and infectious pleural injury, respectively.

**Figure 6.**
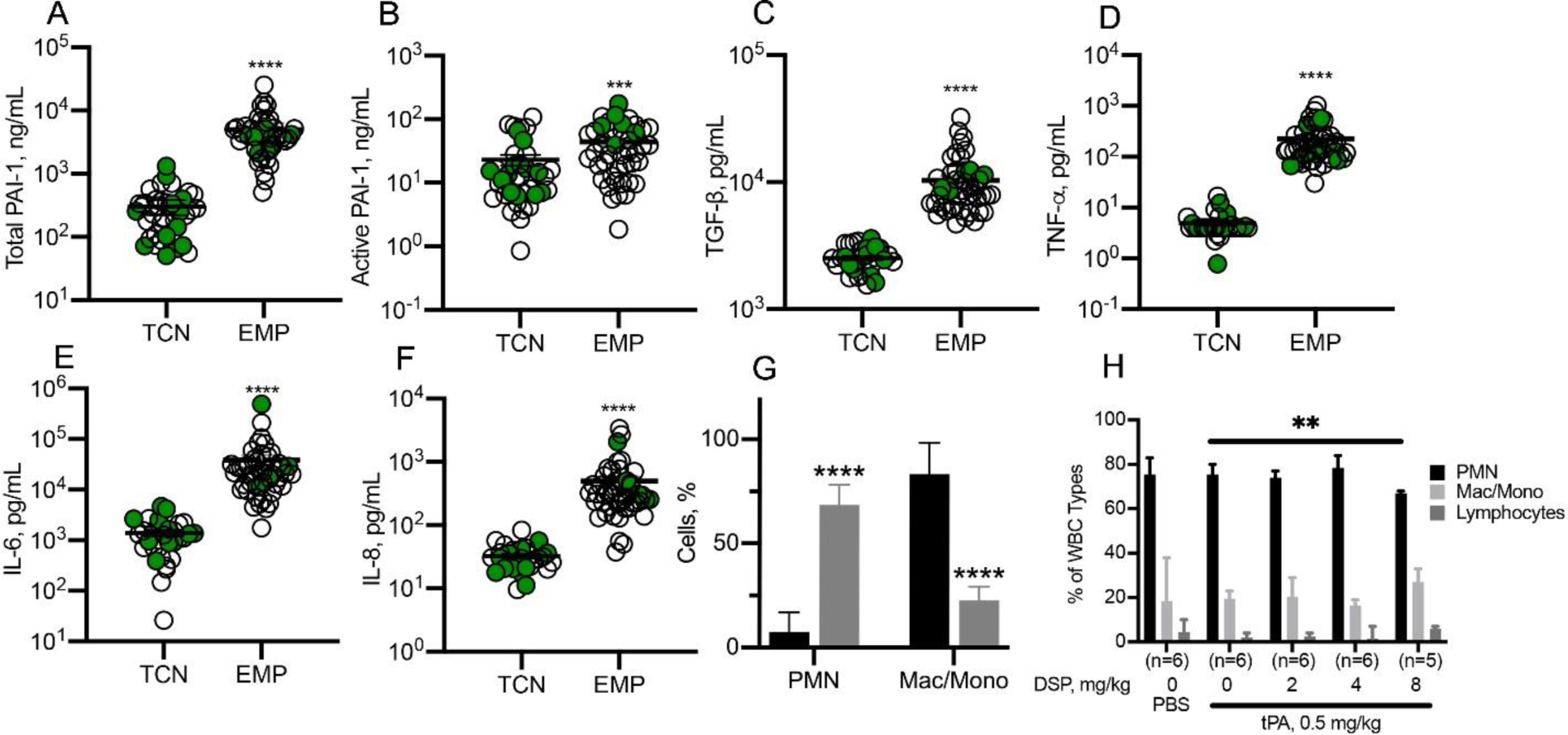
Inflammatory parameters were higher in rabbits with *S. pneumoniae*-induced pleural injury 24h after PAI-1-TFT. Pleural fluid from rabbits with TCN-induced injury or *S. pneumoniae*-induced pleural injury (EMP) were subjected to ELISA to measure: total PAI-1 (**A**), active PAI-1(**B**), TGF-β (**C**), TNF-α (**D**), IL-6 (**E**), and IL-8 (**F**), 24h after PAI-1-TFT. Significantly elevated levels of all targets were detected in pleural fluids of rabbits with empyema. There was no statistically significant difference between different treatments of each type of pleural injury. Data are presented as the mean ± SEM. Significance was determined using an unpaired, 2-tailed Kolmogorov-Smirnov test. \*\*\*\**P*<0.0001 (total PAI-1 n=41, n=45; active PAI-1; n=37, n=44; TGF-β, n=31, n=47; TNF-α, n=31, n=50; IL-6, n=31, n=51; IL-8, n=32, n=49 for TCN-induced injury and empyema, respectively). WBC differential counts in pleural fluids from rabbits with TCN (black) - and *S. pneumoniae*-(grey) induced pleural injury (**D**). Pleural fluids were collected at necropsy. Neutrophil predominance was noted in *S. pneumoniae*-induced injury while a mixed myeloid cell distribution was observed in chemically-induced injury. Results of a Mann-Whitney test on ranks showed a statistically significant change (p<0.0001) in cell populations. Data are presented as a grouped vertical bar graph with standard deviation. Pleural fluid WBC differentials in *S. pneumoniae*-induced PAI-1-TFT treated pleural injury in rabbits (**H**). A significant decrease (*P*<0.01 (**)) in the number of PMN is noted between rabbits treated with 0.5 mg/kg sctPA and 0.5 mg/kg sctPA with 8.0 mg/kg DSP. Significance was determined using an unpaired, 2-tailed Kolmogorov-Smirnov test. Data are presented as a grouped vertical bar graph with standard deviation.

**Figure 7.**
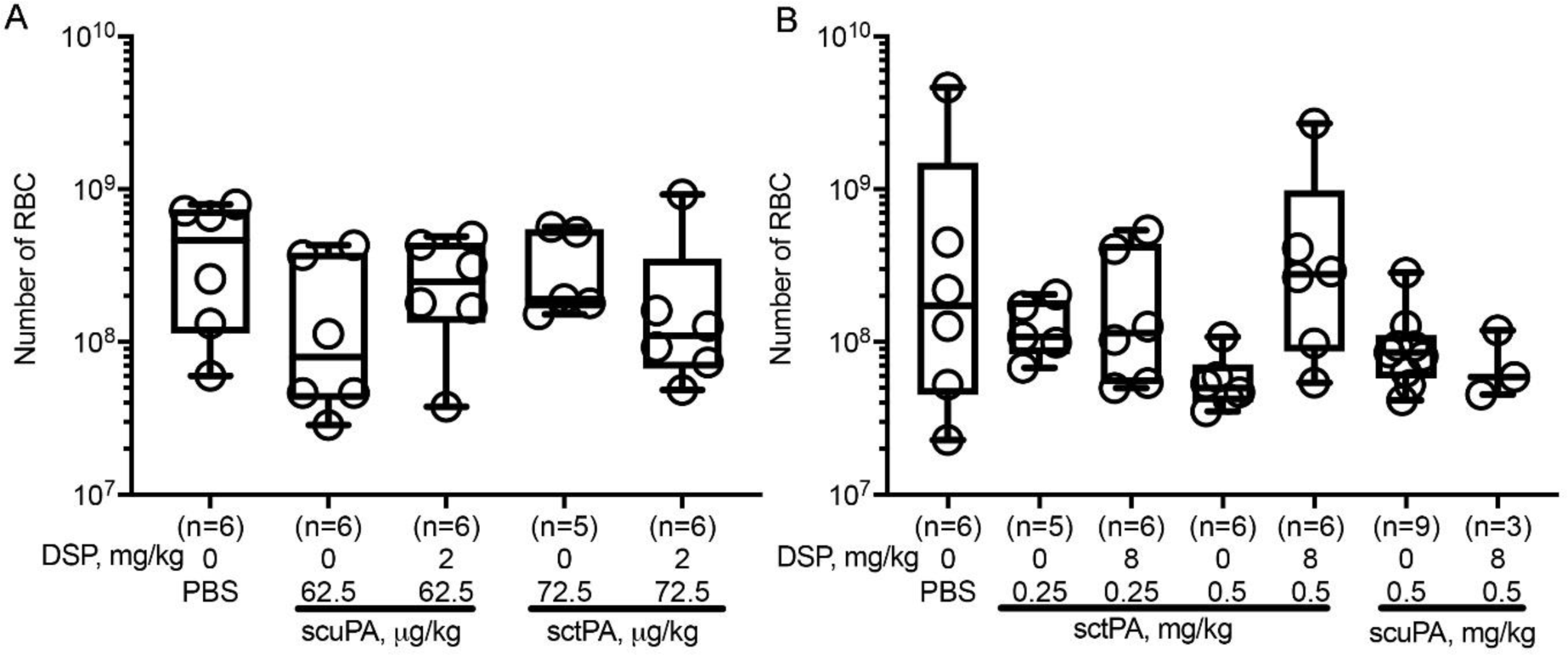
Low dose PAI-1-TFT induces no intrapleural bleeding. Pleural fluids from rabbits with TCN-induced (**A**) or *S. pneumoniae*-induced (**B**) pleural injury were collected at necropsy and RBCs were counted using a Beckman Coulter. **(A)** Animals with TCN-induced pleural injury were treated (from left to right) with: vehicle control (n=6); 62.5 µg/kg of scuPA alone (n=6); and with 2.0 mg/kg DSP (n=6); 72.5 µg/kg of sctPA alone (n=5), and with 2.0 mg/kg DSP (n=6). **(B)** Animals with infectious pleural injury were treated (from left to right) with: vehicle control (n=6); 0.25 mg/kg of sctPA alone (n=5), and with 8.0 mg/kg DSP (n=6), 0.5 mg/kg of sctPA alone (n=6), and with 8.0 mg/kg DSP (n=6), 0.5 mg/kg of scuPA alone (n=9), and with 8.0 mg/kg DSP (n=3). No statistically significant differences in RBCs were observed in either model after treatment with PAI-1-TFT as compared to vehicle treated animals. Data are presented as box and whisker plots (all points shown).

## Discussion

PAI-1, which is elevated in the pleural space by up to three orders of magnitude in empyema (4-6), could be a biomarker of the severity of infectious pleural injury (6). Although identified as a major endogenous, mechanism-based inhibitor of tPA and uPA, to the best of our knowledge, PAI-1 is not considered a molecular target in fibrinolytic therapy of empyema, possibly because an initial multifold excess of the exogenous plasminogen activator is employed. Nevertheless, the combination of rapid clearance of the drug from the pleural space with constant replenishing of PAI-1 due to local overexpression dramatically affects the half-life of intrapleural plasminogen activating activity and fibrinolysis. Low-dose, local PAI-1-TFT allows a significant decrease in the effective therapeutic dose of the drug, minimizing the risk of bleeding complications, and, thereby, expanding the population of patients with empyema that could tolerate pharmacological treatment. The goals of the present study were to validate PAI-1 as a molecular target in two rabbit models of pleural injury (chemical and infectious) that differ in levels of intrapleural PAI-1 (Figure 1). To target the PAI-1 mechanism, we employed DSP, a peptide that interferes with exosite interactions between PAI-1 and plasminogen activator (10;11), thereby competing with PAI-1 for enzymes (15;16). Choosing an animal model that adequately recapitulates empyema in humans is critical for translating the results of PAI-1-TFT to clinical trials. Unlike mice, rabbis possess a fibrin structure similar to humans, and the biochemistry of rabbit PAI-1 and other fibrinolytic enzymes and inhibitors is likewise similar (20-24). Rabbit models allow for testing of human plasminogen activators and have previously been applied successfully in drug development, including the current MIST2 protocol (25) for the treatment of empyema (26;27). Here, we demonstrated that DSP mediated PAI-1-TFT was effective (i) for both sctPA and scuPA in the treatment of TCN-induced pleural injury and (ii) for sctPA but not scuPA in infectious pleural injury in rabbits. In both models, the addition of DSP to fibrinolytic therapy rendered low and otherwise ineffective doses of plasminogen activators effective (Figures 2, 3). Notably, the maximal decrease in the MED of plasminogen activators due to targeting the PAI-1 mechanism with DSP was 8-fold in both models, which was similar to the effect of mAbs observed previously in chemically-induced pleural injury (17). The role of PAI-1 as a molecular target for fibrinolytic therapy in pleural injury is also supported by our previous observations of a decrease in the efficacy of fibrinolytic therapy due to intrapleural stabilization of the active conformation of PAI-1 (14) in a “molecular sandwich” type complex (28). Likewise, an increase in the intrapleural level of PAI-1 in empyema compared to chemically-induced pleural injury (Figure 1) predictably results in an increase in the MED of sctPA and scuPA or failure of therapy with doses previously identified as effective in TCN-induced pleural injury (8). To achieve efficient PAI-1-TFT in empyema, the dose of DSP was also increased from 2.0 mg/kg, which was effective in TCN-induced pleural injury (Figure 2A), to 8.0 mg/kg (Figures 2B, 3A). While DSP (8.0 mg/kg) converted ineffective doses of sctPA (0.5 and 0.25 mg/kg) to effective doses (Figures 2B, 3A), it did not improve the outcomes of treatment with scuPA in empyema (Figure 2A). The striking difference in the effects of DSP on fibrinolytic therapy with scuPA in chemically-induced (Figure 2A) and infectious pleural injury (Figure 2A) suggest differences in the mechanisms of uPA-supported intrapleural fibrinolysis in the two models. The results of LC MS/MS analyses (Figure 4) also demonstrate that an increase in the dose of the adjunct from 2.0 mg/kg results in an increase in the DSP levels in the pleural fluid until PAI-1-TFT becomes effective at 8.0 mg/kg DSP (Figure 4, green symbols). Interestingly, DSP alone shows a trend towards improving therapeutic outcomes in both the chemically-induced and infectious pleural injury models (Figure 2A), indicating a possible effect on the endogenous fibrinolytic system. Targeting the PAI-1 mechanism with DSP differs from PAI-1-TFT with anti-PAI-1 mAbs (14;17), as DSP interacts with the enzyme rather than PAI-1 itself. Thus, DSP could be combined with anti-PAI-1 mAbs to achieve a further increase in the efficacy of PAI-1-TFT due to possible additivity in the effects of different intermolecular mechanisms of modulation of PAI-1 activity and destabilization of acyl-enzymes by DSP and mAbs. *S. pneumoniae*-induced empyema in rabbits (8) recapitulates key features of the disease in humans, including timing and staging: the progression from an early stage, acute empyema to one that is chronic and accompanied by increasing loculation and pleural thickening. An increase in the intrapleural PAI-1 and cytokines in rabbit empyema, compared to chemically-induced pleural injury coincides with an increase in the severity of pleural injury and MED of plasminogen activator (Figures 1, 6). Likewise, levels of PAI-1, TNF-α, TGF-β, IL-8 in loculated/complicated pleural effusions in humans are higher than those in transudate/free flowing ones (5;6). Although the levels of cytokines differ significantly between chemically-induced and infectious pleural injury in rabbits (Figures 1, 6), PAI-1 is the molecular target shared by both models, and thus could be a molecular target in human empyema as well. The rate of intrapleural inactivation of uPA and tPA in acute empyema was similar to that which was observed in chemically-induced pleural injury (Figure 5). Although the rates of intrapleural fibrinolysis in two models of pleural injury are comparable, differences in the levels of the molecular target affect the outcomes of PAI-1-TFT in both models. In previous studies, we detected high levels of plasminogen activating activity in pleural fluids of patients treated with scuPA at 3h after fibrinolytic therapy, which was completely suppressed at 23h (29). Therefore, intrapleural half-life for plasminogen activators in humans and rabbits are within the same order of magnitude and significantly exceed those reported in the circulation. Thus, an increase in the dose of the plasminogen activator up to 50-100 mg proposed for treatment in advanced empyema (30), could be avoided with low dose PAI-1-TFT. Both humans with empyema (8;17;29) and rabbits with infectious pleural injury (8), (Figure 5) feature inhibition of fibrinolytic activity in pleural fluids collected at baseline and at 23-24h post intervention. Activation of plasminogen in pleural fluids at baseline results in fibrinolytic activity (Figure 5E-H) that varies up to two orders of magnitude in human samples (8;17). This observation provides support for the concept of fibrinolytic potential (8;17;29), which characterizes the fibrinolytic system of an individual patient that correlates with therapeutic outcomes in humans treated with scuPA (29). Correlation between fibrinolytic potential at the baseline and at 24h (Figure 5J,L) further supports our hypothesis of a personalized biomarker of an individual’s fibrinolytic system. Notably, the level of fibrinolytic activity in pleural fluids collected 10 min or later after treatment (Figure 5E-H) is significantly lower than that observed in the activated baseline pleural fluids collected just prior to injection of plasminogen activator or at 24h. This could reflect a depletion of pleural fluid plasmin through its binding to intrapleural fibrin, thus increasing the probability of favorable outcomes in individuals with higher fibrinolytic potential (Figure 5J,L; (29)). Likewise, a higher steady state (10-40 min) fibrinolytic activity in pleural fluids from animals treated with scuPA based PAI-1-TFT (Figure 5E,G) compared to that for sctPA (Figure 5F,H) could indicate less plasmin associated with fibrin and contribute to worse outcomes with scuPA in rabbit empyema (Figure 2). In the injured rabbit (14;17;18;31) and human (29) pleural space, uPA forms “molecular cage” type complexes with endogenous αM, which feature a 10-fold longer intrapleural half-life time than free uPA, and could contribute to plasminogen activating activity and fibrininolysis (17;18). Notably, high levels of αM/uPA were detected during PAI-1-TFT of both infectious and chemically-induced pleural injury and were similar to those observed previously (18). Surgical interventions are often offered to the majority of patients with advanced stage empyema. However, a significant proportion of patients are poor candidates for surgery due to comorbidities and advanced age. Since conventional intrapleural fibrinolytic therapy confers a risk of bleeding, fluid drainage using negative pressure remains the only option for this group, which has a high mortality rate (32;33). Our results (Figure 7) clearly indicate that local, low-dose PAI-1-TFT in two models of rabbit pleural injury does not statistically affect the number of RBCs in pleural fluids and, thus, does not increase the risk of intrapleural bleeding complications. Current trends in pharmacological treatment of empyema in humans include (i) de-escalation of sctPA dose to use in combination with deoxyribonuclease (34;35), (ii) evaluation of need of deoxyribonuclease and dose timing when combined with sctPA (36-41) and (iii) comparison of the efficacy of tPA and uPA (29;42-45). Our studies demonstrate that the level of PAI-1 and fibrinolytic potential in pleural fluids (8;17;29) are significant contributors to the outcome of fibrinolytic therapy following pleural injury in two rabbit models. Both parameters could vary up to two orders of magnitude in pleural fluids of human patients with empyema (5;17), revealing an opportunity to personalize fibrinolytic therapy. Notably, with use of low dose PAI-1-TFT, fibrinolytic potential could become a significant predictor of therapeutic success in patients with empyema. In summary, the similarities between human patients with empyema and our model of empyema in rabbits, raises the possibility of eventual translation of PAI-1-TFT to clinical practice. Low-dose PAI-1-TFT was effective and well-tolerated in two models of pleural injury in rabbits and there was no evidence of intrapleural bleeding, the most serious and commonly encountered complication of intrapleural fibrinolytic therapy of empyema. We envision that the PAI-1-TFT therapy presented here could be an effective therapeutic option for a broader population of patients and is safer for those with comorbidities, increased risk of bleeding, or who are otherwise poor candidates for surgical intervention.

## Methods

### Animal Protocols

The animal protocols detailed here were approved by the Institutional Animal Care and Use Committee at The University of Texas Health Science Center at Tyler. Studies used female, pathogen-free New Zealand white rabbits (3.0-3.6kg; average age 18 weeks) from Charles River Laboratories (Wilmington, MA). These studies required a total of 107 rabbits, 43 for the model of TCN-induced pleural injury and 64 for the model of empyema. A model of chemically-induced pleural injury in rabbits was implemented as detailed previously (9). Briefly, a 3 ml mixture of TCN in ascorbic acid (20mg/mL TCN (T3258, Sigma-Aldrich), 2.5 mg/mL ascorbic acid (A7506, Sigma-Aldrich) and lidocaine (1 mg/mL, UTHCT Compounding Pharmacy), was delivered by intrapleural injection. A model of acute empyema in rabbits was used, as described previously (8). Briefly, 1-5 × 10^8^ cfu of *S. pneumoniae* (D39 strain, National Collection of Type Cultures, Salisbury UK) in 3 ml of 0.5% brain-heart infusion agar (BD 238400, BD Diagnostic Systems) was delivered by intrapleural injection. Clavomox (10 mg/kg, subcutaneous, daily for 1-3 days as clinically indicated by Attending Veterinarian) (10000485, Zoetis) was started at 28-30h post-infection. Accumulation of pleural fluid and fibrin deposition was monitored using ultrasonography; detailed in the ultrasonography subsection. After 48 or 72h for TCN-induced pleural injury and empyema, respectively, intrapleural therapeutics were administered depending on treatment group via an 18-gauge catheter, which was flushed with 0.5 ml PBS. In all experiments, samples of pleural fluids (0.5 ml) from each rabbit were collected prior to and at 10, 20, and 40 min after PAI-1-TFT and processed for later analysis (7;8;17;18). Anesthesia, postoperative pain medication, and animal care were provided as reported previously (7;8;17;18). with details provided in the supplement. Animals were monitored carefully for signs of distress, pain, and worsening clinical status. Euthanasia was accomplished using intravenous injection of 1 ml of commercial euthanasia solution (sodium pentobarbital 390 mg/mL and phenytoin 50 mg/mL) followed by exsanguination.

### Ultrasonography

Development of pleural injury was monitored via ultrasonography. Briefly, B-mode ultrasonography of the chest was performed (18;19) using a Logiq e system (GE Healthcare, Milwaukee, WI) using R5.2.x software and a multifrequency transducer model 12L-RS at a working frequency of 10 MHz, as previously described (8).

### Metrics of pleural injury

For each animal, gross lung injury scores (GLIS) were calculated as previously described (7;17;18). PAI-1-TFT was considered successful when GLIS≤10. Multiple fibrin webs or sheets connecting the visceral and parietal pleura or “too numerous to count” (TNTC) strands and a fibrinous lung coating correlated with a GLIS = 50. Pleural thickening was measured using morphometry, as previously described (8).

### DSP

DSP (EEIIMD) was synthesized by GenScript.

### Pharmacokinetic analysis

The residual DSP in pleural fluid samples was determined using LC MS/MS. Briefly, collected samples were extracted using acetonitrile, cleaned with Microcon centrifugal filters (MRCPRT010, MilliporeSigma) and injected via autosampler onto a Pinnacle DB C8 Column (9413332, Restek) reverse-phase column operated at a flow rate of 200 μl/min. A gradient elution method using solvent A (99% H2O, 1% acetonitrile, 0.05% TFA) for 5 min, a gradient to 100% solvent B (1% H2O, 99% acetonitrile, 0.05% TFA) over 25 min, and 100% solvent B for 2 min was employed. Mass spectrometric analyses were performed on a TSQ Vintage (Thermo Fisher Scientific) mass spectrometer equipped with an electrospray ion source operating in positive mode and SRM detection mode. The apparent rate constant of intrapleural elimination of DSP (k_clr_) was estimated by fitting a single exponential equation to the plot of concentration of DSP in pleural fluids on time.

### ELISA for quantitation of antigens and PAI-1 activity

Commercial ELISA were used to assess the levels of total and active PAI-1 (RBPAIKT-TOT, RbPAIKT; Molecular Innovation Inc.), TNF-α (DY5670, R&D Systems), TGF-β (DY240-05, R&D Systems), IL-6 (DY7984, R&D Systems), IL8 (ELL-IL8-1, RayBiotech) in rabbit pleural fluids.

### Amidolytic, plasminogen activating and fibrinolytic activity assays

Amidolytic (18;31), plasminogen activating (18;31), and fibrinolytic activity (17;46), and concentration of αM/uPA complexes (18;31) in pleural fluids were measured as previously described.

### Statistics

Statistical significance was determined using the Kruskal-Wallis test as well as pairwise multiple-comparison procedures (Dunn’s multiple comparison test). Correlation coefficients (*r*) were calculated using the fit of the curve and used as a parameter to determine the best fit. Data were analyzed using SigmaPlot 12.3 for Windows (SPSS Inc.) and GraphPad Prism 8.2.1, as previously described (8;17;18).

### Study approval

All animal procedures were approved by the Institutional Animal Care and Use Committee at The University of Texas Health Science Center at Tyler. Studies used female, pathogen-free New Zealand white rabbits (3.0-3.6kg; average age 18 weeks) from Charles River Laboratories (Wilmington, MA). (IACUC protocols 499,616, 617)

## Author Contributions

GF, RG, KS, SK, CJDV, DM, MC, KK, AAK performed experiments and data collection. AOA and AB assisted in animal experiments. AAK, RG, GF analyzed, plotted and interpreted the data. AAK, RG and GF performed statistical analyses. SK edited the manuscript. SI and DBC revised the manuscript. AAK conceived the study, interpreted the data and wrote the manuscript which was reviewed, and approved by all authors.

## Acknowledgments

The authors thank James Henry and vivarium staff of The University of Texas Health Science Center at Tyler for excellent technical support and care for animals. This work was supported by NIH grant R01HL130402 (GF, SI, AAK) and The Texas Lung Injury Institute.

